# Nanobodies reveal an extra-synaptic population of SNAP-25 and Syntaxin 1A in hippocampal neurons

**DOI:** 10.1101/472704

**Authors:** Manuel Maidorn, Aurélien Olichon, Silvio O. Rizzoli, Felipe Opazo

## Abstract

Synaptic vesicle fusion (exocytosis) is a precisely regulated process that entails the formation of SNARE complexes between the vesicle protein synaptobrevin 2 (VAMP2) and the plasma membrane proteins Syntaxin 1 and SNAP-25. The sub-cellular localization of the latter two molecules remains unclear, although they have been the subject of many recent investigations. To address this, we generated two novel camelid single domain antibodies (nanobodies) specifically binding to SNAP-25 and Syntaxin 1A. These probes penetrated more easily into samples and detected their targets more efficiently than conventional antibodies in crowded regions. When investigated by super-resolution imaging, the nanobodies revealed substantial extra-synaptic populations for both SNAP-25 and Syntaxin 1A, which were poorly detected by antibodies. Moreover, extra-synaptic Syntaxin 1A molecules were recruited to synapses during stimulation, suggesting that these are physiologically-active molecules. We conclude that nanobodies are able to reveal qualitatively and quantitatively different organization patterns, when compared to conventional antibodies.

## Introduction

The release of neurotransmitter by synaptic vesicle fusion is an extremely rapid process, which follows neuronal stimulation with high precision. Its control relies on several proteins that serve to dock synaptic vesicles at the fusion site (active zone) or to sense increases in the intracellular Ca^2+^ concentration, which marks the physiological trigger for fusion.^1^ The act of fusion itself, however, relies almost exclusively on three soluble N-ethylmaleimide sensitive factor attachment receptor (SNARE) proteins: Synaptobrevin 2 (VAMP2), located on synaptic vesicles, and the plasma membrane proteins Syntaxin 1A and SNAP-25.^2^ They mediate exocytosis by forming a heteromeric helical bundle that brings the two membranes together and forces their fusion.^3^ All of these molecules are highly abundant in synaptic terminals, with ~70 VAMP2 molecules on average found on the synaptic vesicle^4, 5^ and with thousands of SNAP-25 and Syntaxin 1A molecules on the synapse surface.^5^

While the localization of synaptobrevin 2 on the vesicle surface is fairly clear, the location of SNAP-25 and Syntaxin 1A is less understood. Investigation by super-resolution imaging revealed that both SNAP-25 and Syntaxin 1A are enriched at synapses.^5^ At the same time, analyses by immuno-electron microscopy have provided less clear-cut results, with the proteins present both in synapses and in other axonal regions.^6^ In addition, both SNAP-25 and Syntaxin 1A have been studied in neuroendocrine PC12 cells, where their distributions have been thoroughly characterized. In brief, both molecules form clusters.^7–10^ More specifically, Syntaxin 1A forms dense clusters of ~75 molecules grouped in a roughly circular area with a diameter of 50-60 nm^8^ while SNAP-25 appears to form larger and more loose clusters of ~130 nm in diameter.^9^ Cluster formation does seem to take place also at synapses, with the clusters possibly containing both SNAP-25 and Syntaxin 1A molecules, at least to some extent.^11^ The localization of these proteins has also been investigated by live-cell imaging, using GFP-tagged proteins. While these GFP-tagged molecules showed clusters in PC12 membranes,^10^ a GFP-tagged Syntaxin 1A chimera revealed an overall homogenous labeling pattern on the plasma membrane of cultured neurons.^12^

At the same time, a caveat of most super-resolution studies on membrane proteins has been the application of conventional antibodies for target detection. It was repeatedly demonstrated that the large size of the antibodies (~10-15 nm) may compromise both the resolution and the signal distribution in super-resolution fluorescence microscopy.^13–15^ At the same time, the facts that both primary and secondary antibodies have two epitope-binding domains (bivalency) and that multiple types of secondary antibodies are often used simultaneously (polyclonality), pose significant problems, since they can induce artificial aggregation of antibodies.^16, 17^ This has rendered some of the clustered (spotty) patterns described in super-resolution investigations of immunostaining doubtful^18^ and has encouraged researchers to develop alternative affinity probes. Small probes like aptamers^18^ and nanobodies have already been used to enhance the resolution attained in biological samples when compared to conventional antibodies in super-resolution microscopy,^13, 14^ albeit only a handful are currently available. One prominent example are camelid single domain antibodies (sdAbs), also termed nanobodies, which are derived from antibody types lacking the light chains.^19^ Nanobodies exhibit several properties beneficial for molecular target detection, including small size (~2-3 nm), monovalency, monoclonality and their recombinant production, which allows easy functionalization like site-specific and stoichiometric labeling.^19, 20^

Here we present two novel nanobodies that were selected and produced to detect the synaptic proteins SNAP-25 and Syntaxin 1A with high specificity and affinity. Syntaxin 1A is one of two isoforms of this molecule expressed in the nervous system. These isoforms (1A and 1B) show overlapping, albeit not identical distributions,^21, 22^ similar functions,^23^ and similar levels in central nervous system synapses.^5^ Using nanobodies for SNAP-25 and Syntaxin 1A, we could reproduce some of the previously published results on the distribution of these molecules in neurons, although the staining pattern presented by the nanobody suggested a far smoother staining than the one obtained by antibodies. Interestingly, the nanobodies also revealed large populations of both SNAP-25 and Syntaxin 1A outside the synapses, which were poorly revealed by the antibodies. Furthermore, the extra-synaptic Syntaxin 1A molecules, but not the SNAP-25 molecules, were recruited to the synaptic boutons center upon strong neuronal stimulation. In addition, two-color investigations using super-resolution microscopy also showed that the SNAP-25 and Syntaxin 1A are better correlated than previous antibody-based measurements have suggested both within and outside synapses. Overall, these findings suggest that small, monovalent probes such as nanobodies are able to detect not only quantitative, but also qualitative differences in molecular distribution, when compared to antibodies.

## Results

### Screening and characterization of nanobodies for SNAP-25 and Syntaxin 1A

To obtain nanobodies for SNAP-25 and Syntaxin 1A (Figure 1A), we first immunized an alpaca *(Vicugna pacos)* with rat Syntaxin 1A lacking its C-terminal transmembrane domain (residues 1-262), and with full-length rat SNAP-25. The cysteine residues of the latter were mutated to serines, to facilitate expression in *E. coli* (Figure 1B). After two panning rounds of phage display^24^ (Supplementary Figure 1A), 22 and 11 ELISA-positive families of nanobodies were identified for SNAP-25 and Syntaxin 1A, respectively. The most abundant member present in each of the families was produced in *E. coli* in a small scale and their specificity was further evaluated by dot-blot assays. Such candidates were then used to immunostain fibroblast cells transiently expressing SNAP-25 or Syntaxin 1A fused to EGFP (Supplementary Figure 1B). The nanobodies showing specific signals and minimal background in the immunostaining were subsequently sub-cloned into a bacterial expression vector that includes a cysteine at their C-terminus for direct conjugation to a fluorophore.^25^

**Figure 1:**
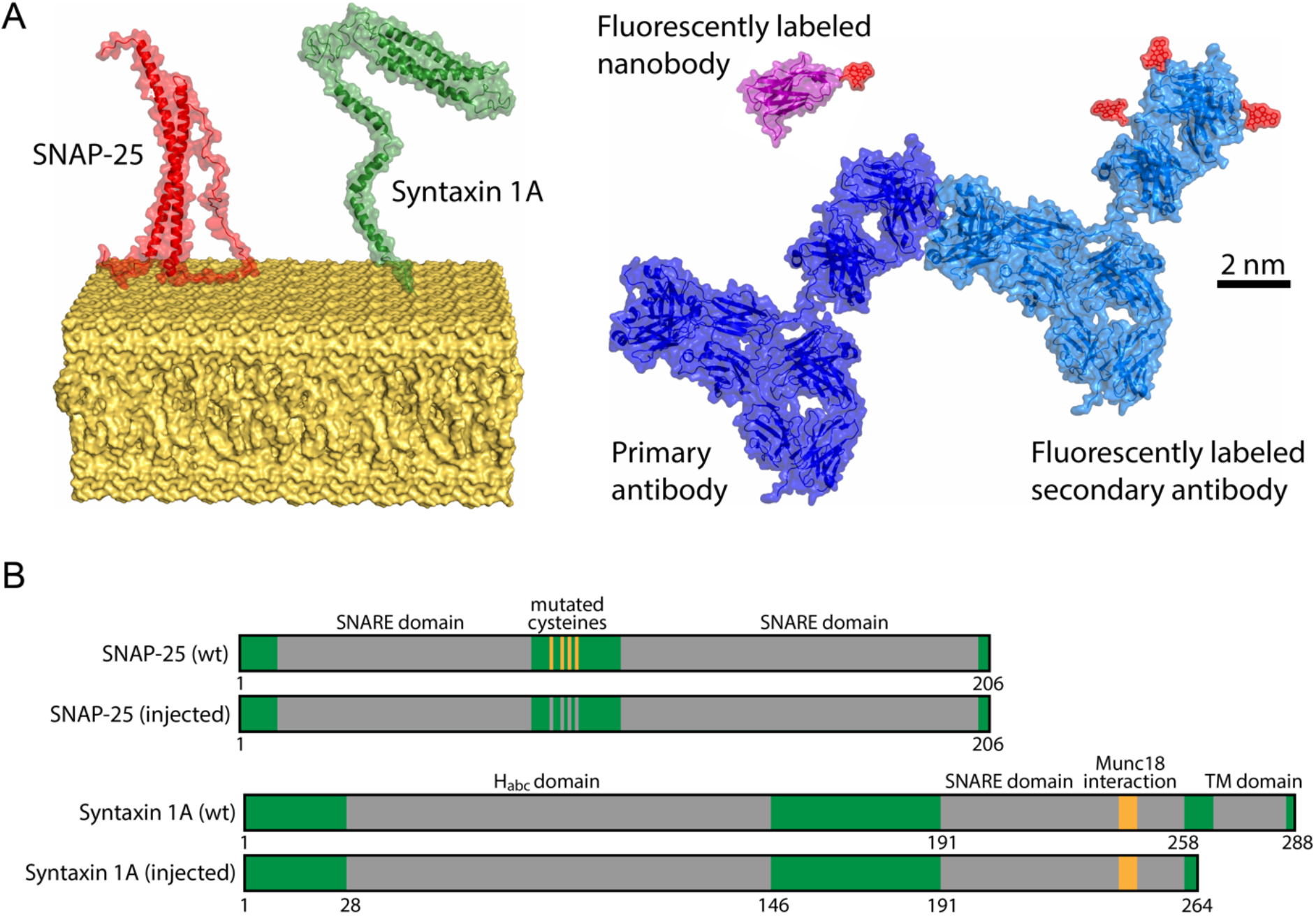
Schematics of the proteins involved in this study. **(A)** Molecular models of SNAP-25 (based on PDB: 1KIL, in red) and Syntaxin 1A (based on PDBs: 1HVV & 1BR0, in green) residing in the plasma membrane (in yellow). On the right we show a nanobody (PDB: 1I3V, in purple) bearing a single Atto647N at the C-terminus, and a complex of a primary and a randomly-labeled secondary antibody with Atto647N on lysines (PDB: 1IGY in blue and light-blue). All molecular models are displayed in the same scale and the bar represents 2 nm. **(B)** Schematic view of antigens used for the immunization and their wild-type forms. Important functional domains are marked in gray (TM=trans-membrane domain) and the amino acids positions are denoted below. For immunization (injected), all four cysteine residues of SNAP-25 were mutated to serines. For injection of Syntaxin 1A, only the cytosolic portion of the molecule was used to facilitate it expression in *E. coli.*

The nanobody candidates termed **S25-Nb10** and **Stx1A-Nb6** performed best for the immunostainings of SNAP-25 and Syntaxin 1A, respectively, and were used for all further experiments. As a first step, dissociation constants (K_D_) were determined by microscale thermophoresis. We found that S25-Nb10 binds to recombinant SNAP-25 with a K_D_ of 15.5±3.3 nM, and that Stx1A-Nb6 binds to recombinant Syntaxin 1A with a K_D_ of 5.0±1.2 nM *in vitro* at room temperature (Supplementary Figure 2). Monovalent probes with dissociation constants in this range are considered high affinity binders.^26^

To identify the epitopes of the nanobodies, we tested different truncated constructs of both SNAP-25 and Syntaxin1A for nanobody binding. Equimolar amounts of these constructs were spotted on a nitrocellulose membrane and were detected by the respective fluorescently labeled nanobodies in dot-blot assays (Figure 2). The blots suggested that S25-Nb10 binds within the first 86 N-terminal residues of SNAP-25, which is one of the two alpha helixes that SNAP-25 contributes to a SNARE complex.^27^ Stx1A-Nb6 binds within the first 112 residues of the N-terminal portion of Syntaxin 1A, which are part of the regulatory Habc domain of Syntaxin 1A.^28^

**Figure 2:**
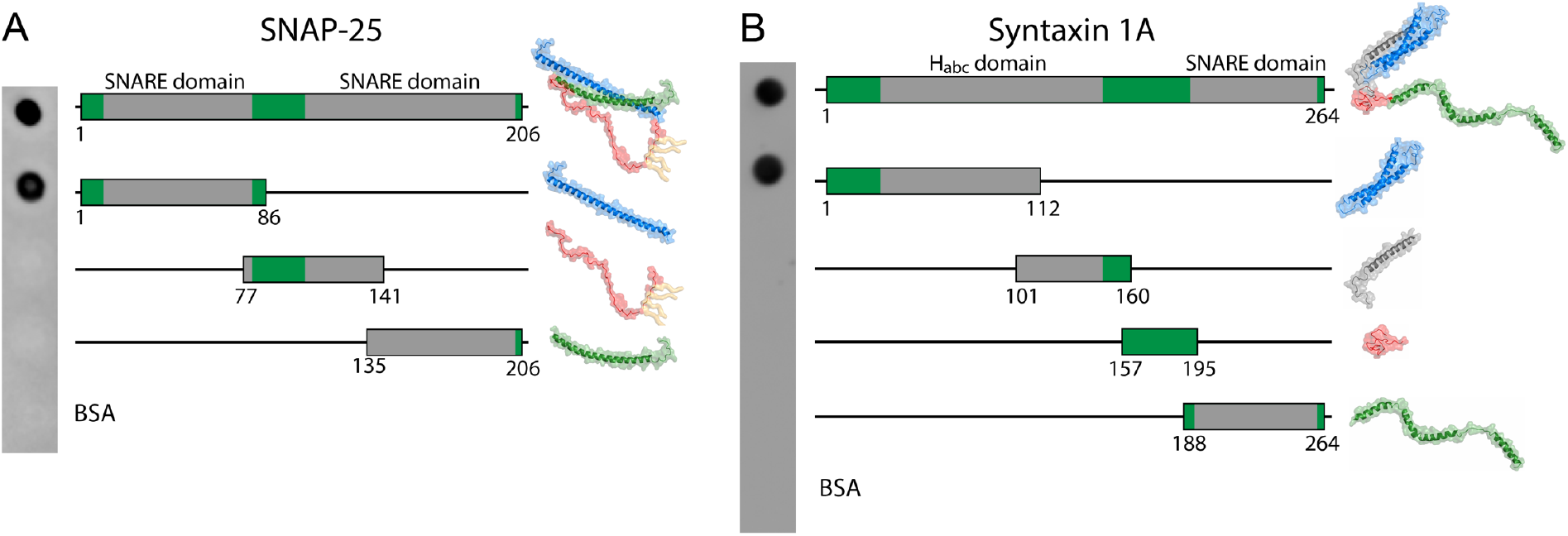
Rough mapping of the binding epitopes of the selected nanobodies. Full length antigen or truncated versions were produced in *E. coli* and were spotted on a nitrocellulose membrane in equimolar amounts. Bovine serum albumin (BSA) was used as negative control. **(A)** S25-Nb10 and **(B)** Stx1A-Nb6 were directly labeled with a single Atto647N fluorophore on their C-terminus and were used for protein detection. The schematics display the location and size of the truncated epitopes (indicated by numbers below). For orientation, molecular structures of the full length and the different truncated domains are displayed nearby.

After determining the binding strength and epitope localization of the nanobodies, we proceeded to evaluate their specificity within their target families. Several homolog proteins are known for both SNAP-25 and Syntaxin 1A (Supplementary Figure 3A, 3B). We therefore tested the binding of the nanobodies to the closest homologs by dot-blot assays. Both nanobodies displayed high specificity, with minimal cross-reactivity to any of the tested homologs (Supplementary Figure 3C, 3D).

We next focused on characterizing the nanobody performance in cellular immunostainings. As a first attempt, we transiently expressed wildtype rat SNAP-25 or Syntaxin 1A, fused to EGFP, in a fibroblast cell line (COS-7 cells), where these proteins are not expressed endogenously. SNAP-25-EGFP was distributed as expected, on the membranes of the transfected cells (Figure 3A), while Syntaxin 1A-EGFP was largely confined to the endoplasmic reticulum, which is typical for cells lacking neuronal binding partners that aid in its transfer to the plasma membrane.^29^ Nevertheless, the cells served as a good first model to test the nanobody immunostaining abilities. They revealed accurately the EGFP-containing cells, and showed no signal in cells not expressing SNAP-25-EGFP or Syntaxin 1A-EGFP (Figure 3A, 3B). The nanobodies recognized specifically their intended targets, whereas SNAP-25 nanobodies did not reveal Syntaxin 1A and *vice versa* (Supplementary Figure 4).

**Figure 3:**
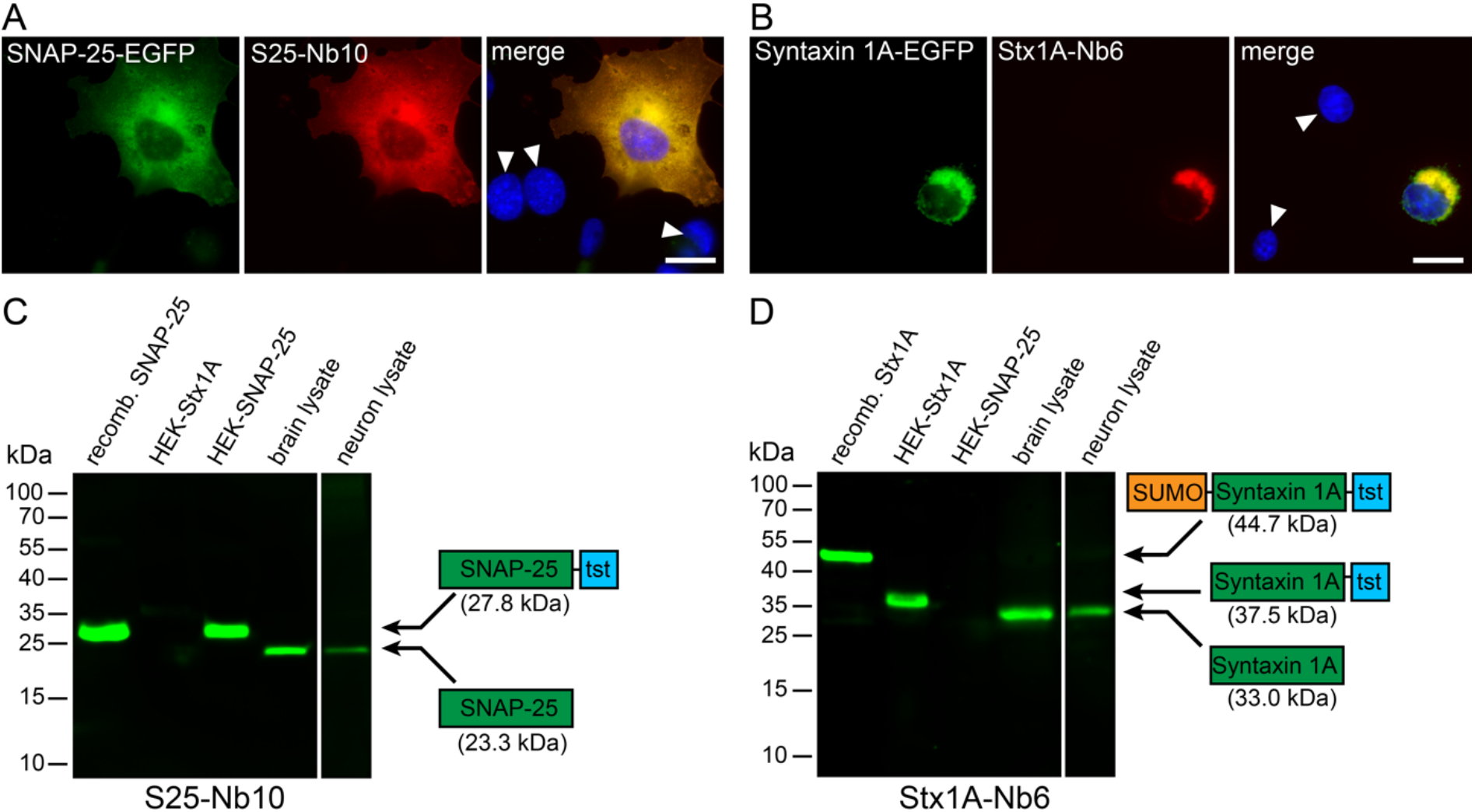
Specificity test for the selected nanobodies in immunofluorescence and Western Blots. **(A & B)** COS-7 cells transiently transfected with SNAP-25-EGFP or Syntaxin 1A-EGFP were stained with S25-Nb10 or Stx1A-Nb6 conjugated with a single Atto647N fluorophore. The nanobody signal correlates with the EGFP signal and shows no staining in untransfected cells (revealed by Hoechst-nuclear staining; white arrowheads). The scale bar represents 10 μm. Note that Syntaxin 1A accumulates in the ER-Golgi region due to an impaired export caused by the lack of neuronal cofactors, as it was found in the past^30^ **(C & D)** The following samples were loaded in a denaturating SDS-PAGE and were blotted on a nitrocellulose membrane (2 μg of each purified protein, and 20 μg of total protein for the cell or brain lysates): *E.* coli-purified full length SNAP-25 including a Twin-Strep-Tag (tst; 27.8 kDa), or Syntaxin 1A fused to a SUMO domain and a Twin-Strep-Tag (44.7 kDa); lysates from HEK293 cells transiently transfected with full length Syntaxin 1A-tst (HEK-Stx1A, 37.5 kDa) or SNAP-25-tst (HEK-SNAP-25, 27.8 kDa); whole rat brain and rat primary hippocampal neurons (15 days *in vitro).* Detection was performed using S25-Nb10 **(C)** or Stx1A-Nb6 **(D)** conjugated to a single Atto647N. Both candidates specifically detect the bands at the expected molecular weights (displayed in the protein schematics), while showing no cross-reactivity to the opposite antigen or to any other proteins present in the lysates.

To complete the nanobody characterization, we also investigated the binding to their targets in a western-blot (WB) assay. For this, we investigated by SDS PAGE the following: Purified recombinant SNAP-25 or Syntaxin 1A produced in *E. coli,* lysates of HEK293 cell lines transiently expressing SNAP-25 or Syntaxin 1A, lysates of total rat brain and lysates of rat primary hippocampal cultures. Both nanobodies were able to bind their corresponding targets accurately (Figure 3C, 3D) and detected no other bands on any of the lanes. Moreover, the bands detected by the nanobodies match those bands detected by commonly used SNAP-25 and Syntaxin 1A antibodies (Supplementary Figure 5A, 5B). We further extended this specificity study by using lysates from mouse liver, muscle, heart, testes and brain. The nanobodies detected no bands in any tissues other than brain, which again suggests that they are both highly specific (Supplementary Figure 5C, 5D).

Finally, we tested whether these nanobodies would be able to bind SNAP-25 and Syntaxin 1A when engaged in a SNARE complex. We conjugated them to maleimide-functionalized agarose beads and aimed to immunoprecipitate SNAP-25 or Syntaxin 1A from whole rat brain lysates (Supplementary Figure 6A, 6B), and from an *in* vitro-formed SNARE complexes (Supplementary Figure 6C, 6D).^3^ Both nanobodies were able to immunoprecipitate their targets, in both experiments. Moreover, they could also co-immunoprecipitate the other SNARE partner, suggesting that S25-Nb10 and Stx1A-Nb6 are able to bind their targets even when they are assembled in SNARE complexes.

### S25-Nb10 and Stx1A-Nb6 detect more epitopes in crowded regions and penetrate more efficiently into thick tissue samples

After the initial characterization of the SNAP-25 and Syntaxin 1A nanobodies, we studied their behavior compared to conventional antibody immunostainings. We first analyzed co-localizations between conventional antibodies and the corresponding nanobodies by confocal laser scanning microscopy in undifferentiated PC12 cells (which have been used extensively to investigate these SNAREs, as indicated in the Introduction). As expected, we observed a strong co-localization between antibodies and nanobodies on PC12 cells using confocal microscopy (Figure 4A), especially at the plasma membranes. However, both nanobodies also showed a distinctive signal in the perinuclear region, where antibodies displayed a poorer detection. As membrane proteins like Syntaxin 1A and SNAP-25 are expected to be present in the ER-Golgi endomembrane traffic system^31^ located in the perinuclear area, we hypothesized that the antibodies may not be able to detect their epitopes efficiently in this tightly crowded region.

**Figure 4:**
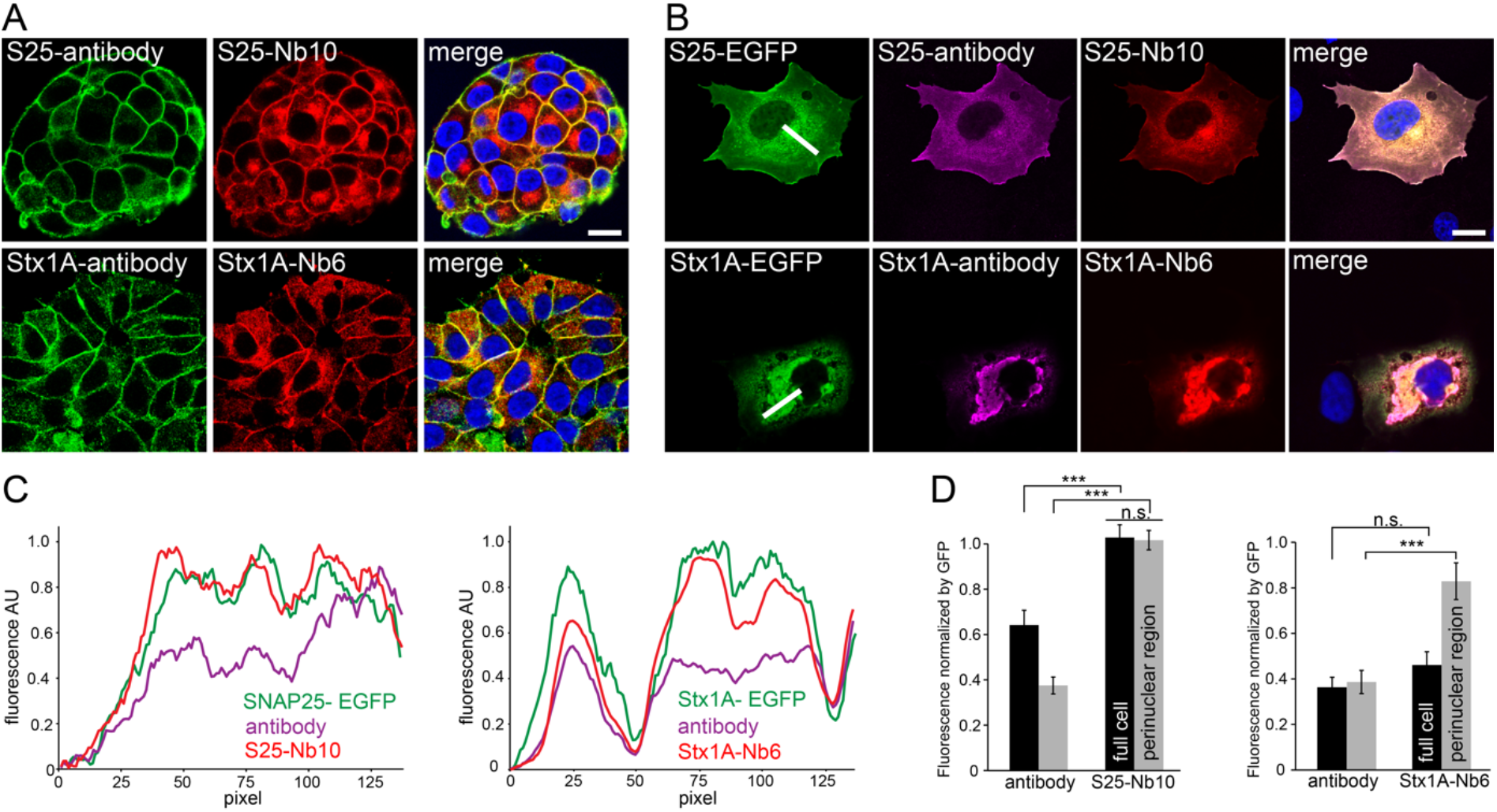
Both nanobodies reveled epitopes that are not reported by antibodies in highly crowded regions of endogenously-expressing PC12 cells or over-expressing COS-7 cells. **(A)** PC12 cells were co-stained with conventional monoclonal antibodies and with our fluorescently labeled nanobodies. Monoclonal anti SNAP-25 and Syntaxin 1A (clone 71.1 and HPC-1) were detected with a secondary antibody conjugated to Abberior-Star580 (in green). S25-Nb10 or Stx1A-Nb6 were conjugated to a single Atto647N fluorophore (in red). Laser scanning confocal images of the nanobody and the antibody signals colocalize relatively good at the plasma membrane, but the nanobodies also reveal stronger signals in the perinuclear areas (especially evident for SNAP-25). The scale bar represents 10 μm. **(B)** Laser scanning confocal images of COS-7 cells transiently transfected with SNAP-25 or Syntaxin 1A fused to EGFP (in green) and co-stained with monoclonal primary and Cy3-fluorescently labeled secondary antibody (clone 71.1 and clone 78.2, in magenta) and nanobodies directly conjugated to Atto647N (in red). Scale bar represents 10 μm. **(C)** Line profiles of the white lines displayed in **(B)** are shown for all three channels, each channel was normalized to its maximum signal intensity on the picture. **(D)** The average fluorescence of full cells, black bars or at the perinuclear regions, light-grey bars in respect to their EGFP signals were calculated. For every selected region of interest, the averaged fluorescence signal of antibodies or nanobodies was calculated and normalized to the average EGFP-fluorescence in the respective region. Statistical analysis was done using unpaired t-test, error bars represent SEM, from three independent experiments analyzing a total of 37 cells for SNAP-25-EGFP and 36 cells for Stx1A-EGFP. n.s. = not significant

To address this hypothesis, we again turned to COS-7 cells and to transient expression of EGFP fusion chimeras of the two SNARE proteins (Figure 4B). We co-immunostained the cells with both antibodies and nanobodies and investigated the correlation of the resulting signals with the EGFP intensity. We found that both nanobodies were able to find more target epitopes in the perinuclear regions, where the EGFP intensity was at its highest (Figure 4B). Exemplary line profiles drawn along these crowded regions revealed that the nanobody signals correlated better with the EGFP signals compared to the respective antibody (Figure 4C, 4D). An analysis of the immunostaining signal intensities relative to the EGFP intensities, revealed that the nanobodies provided brighter signals, especially in the highly crowded perinuclear areas (Figure 4C, 4D). We ruled out epitope competition between the nanobodies and the antibodies to be responsible for this effect, as control experiments did not indicate such an effect (Supplementary Figure 7).

The previous experiments suggested that the antibodies can have difficulties in reaching all epitopes, especially in crowded regions. This problem was also evident when staining tissue samples, rat brain slices of ~35 μm thickness. The nanobodies labeled the slices throughout their entire depth, while the antibodies labeled principally the top and bottom layers, but not the mid-regions of the slices indicating impaired penetration of the antibodies (Supplementary Figure 8).

### Nanobodies reveal an extra-synaptic population of SNAP-25 and Syntaxin 1A in hippocampal cultured neurons

Having broadly characterized the nanobodies, we proceeded to investigate their performance in primary cultured hippocampal neurons (Figure 5 and Supplementary Figure 9A) where both nanobodies did bind their targets abundantly (Figure 5A-5C). To exclude that signals were due to a non-specific interaction between the fluorophore conjugated to the nanobodies and the neuronal membranes,^32^ we also immunostained neuronal cultures with a nanobody directed against GFP, conjugated to the same fluorophore (Atto647N), which displayed no substantial signal under the same conditions (Supplementary Figure 9B).

**Figure 5:**
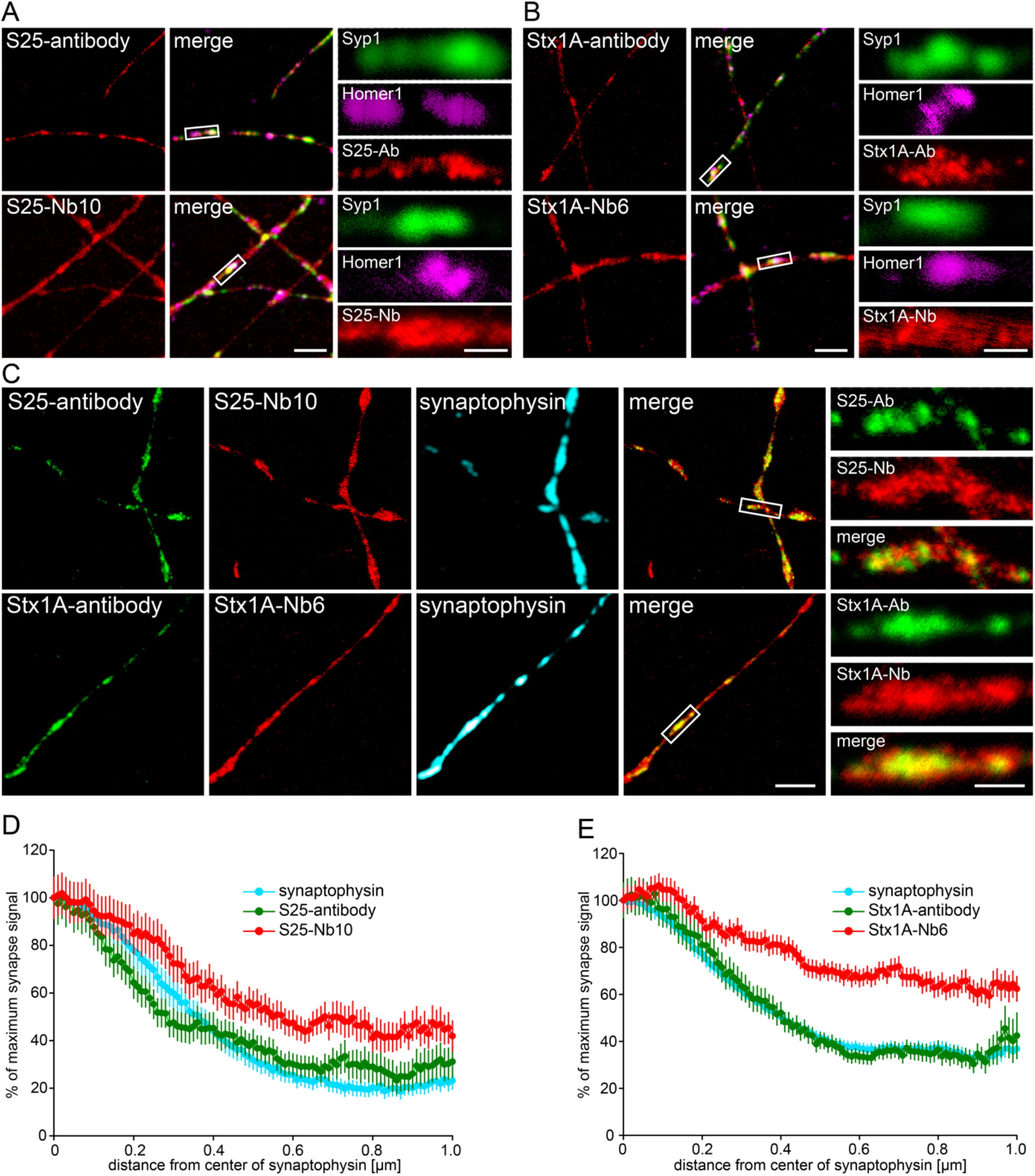
STED microscopy shows that nanobodies detect an extra-synaptic population of SNAP-25 and Syntaxin 1A in primary hippocampal neurons. **(A & B)** Cultured neurons (14-16 days *in vitro)* were co-stained with SNAP-25 or Syntaxin 1A monoclonal antibodies (clone 71.1 or 78.2, respectively) or with the nanobodies bearing Atto647N fluorophores (in red) and with the pre-synaptic marker antibodies against Synaptophysin 1 (Syp1) and the post-synaptic marker Homer1. The merge panels display the synaptic markers and the other co-stained protein in red. Zoomed region of synapses suggest that antibodies and nanobodies are enriched at synapses, but the signals from the nanobodies are also present in extra-synaptic areas. The scale bars represent 2 μm and 500 nm in the low and high zoom, respectively. **(C)** For two-color STED microscopy, primary neurons were co-stained using the same monoclonal antibodies as in (A), detected using a secondary antibody conjugated to Abberior-Star580 and with the nanobodies coupled to Atto647N. Synapses were located in confocal mode with Synaptophysin antibodies as in **(A)**. The merge panel only includes the two STED channels for simplicity. The zoomed areas show each channel and the overlap of both antibody and nanobody STED signals. The scale bars represent 2 μm and 500 nm in the low and high zoom, respectively. **(D & E)** Analysis of the signal distribution up to 1 μm from the center of a synapse (determined by the center of mass of the Synaptophysin staining). We analyzed 176 synapse line scans from six independent experiments for SNAP-25 and 632 synapse line scans from six independent experiments for Syntaxin 1A. The nanobody signals are significantly higher than the antibody signals outside synapses for both experiments (Wilcoxon rank sum tests; p = 0.0016 for SNAP-25; p < 0.00001 for Syntaxin 1A).

We then analyzed the samples by two-color Stimulated Emission Depletion (STED) microscopy. This revealed a different pattern of the target protein distribution between the antibodies and the nanobodies. The antibodies showed a spotty pattern, concentrated mainly in synapses that were revealed by immunostaining for the synaptic vesicle marker Synaptophysin.^4^ The nanobodies also revealed a significant signal in the surrounding areas, suggesting that substantial populations of these proteins are also present outside synapses.

To analyze this observation, we located the center of mass of the Synaptophysin signals, taken as the centers of synapses. We then performed line profiles along the axons for up to 1 μm, starting from the synapse centers (Figure 5D, 5E). The results confirmed that both nanobodies and antibodies reveal more target molecules within synapses (enriched at the synapses), but the nanobodies also revealed a significantly higher population of these molecules outside of synapses when compared to the antibodies. Several explanations can be found to interpret this difference (see Discussion section). Notably, the staining obtained with nanobodies resembles more closely the signals described in the literature with GFP-tagged molecules in living neurons.^12^

One relatively puzzling observation derived from co-immunostainings of SNAP-25 and Syntaxin 1A in neurons has been that these two molecules did not seem to overlap greatly when investigated by super-resolution microscopy.^11^ This was also the case in our hands, with SNAP-25 and Syntaxin 1A clusters appearing to only roughly correlate in axons when using antibodies (Figure 6A). The signal correlation was much stronger when using nanobodies compared to antibodies (Figure 6A, 6B). In spite of the separate signals obtained in antibody stainings, this observation suggests that the two molecules may behave in a relatively similar fashion and may not cluster in widely different regions.

**Figure 6:**
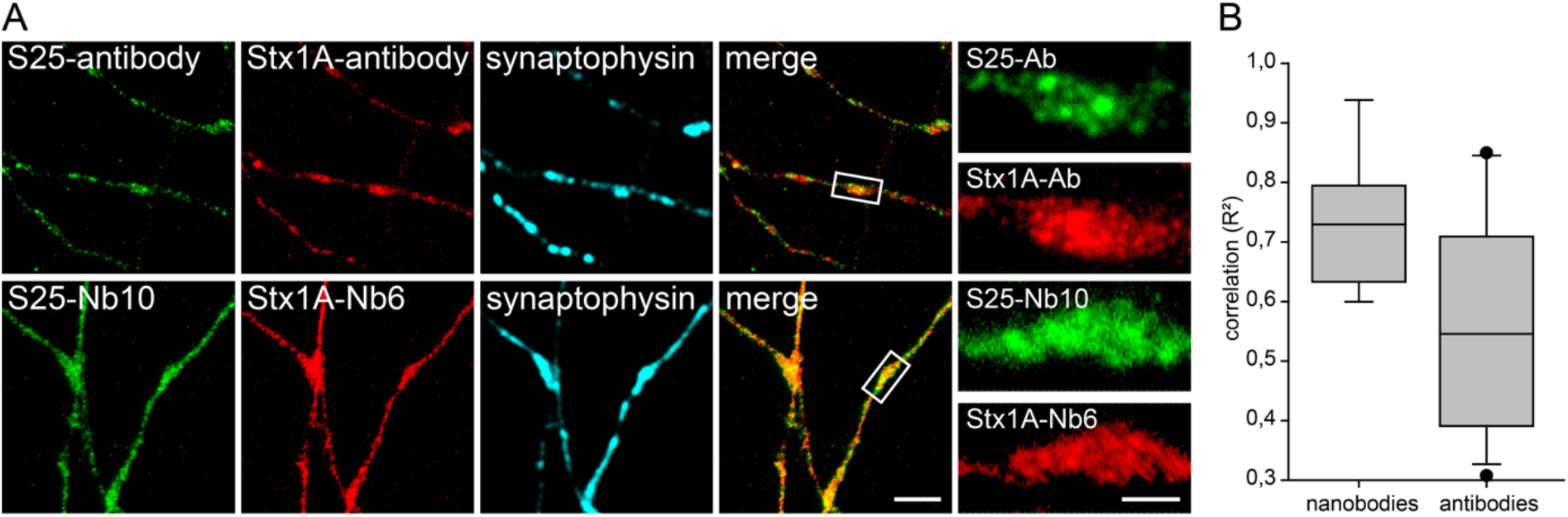
Co-localization between SNAP-25 and Syntaxin 1A, detected by antibodies or nanobodies in STED microscopy. **(A)** Co-staining for SNAP-25 and Syntaxin 1A in primary hippocampal neurons. A polyclonal rabbit antibody against SNAP-25 and a mouse monoclonal against Syntaxin 1A were used. Rabbit anti SNAP-25 was further detected with a secondary antibody conjugated to Abberior-Star580; Syntaxin 1A primary antibody was detected with an Atto647N-conjugated secondary antibody. Nanobody co-staining was performed with S25-Nb10 conjugated to Abberior-Star580 and with Stx1A-Nb6 conjugated to Atto647N. The scale bars represent 2 μm and 500 nm in the low and high zoom, respectively. **(B)** Pearson’s correlation coefficients (expressed as coefficients of determination, R^2^) between the green and red signals within synapses were calculated from 17 independent experiments for the antibodies and from 9 independent experiments for the nanobodies (typically, 10 images per experiment were analyzed). The values are shown as box plots, with the median, 25^th^ and 95^th^ percentile shown in the graph, and with symbols showing outliers. The difference is significant (Wilcoxon rank sum test; p = 0.0311).

To formally characterize these clusters, we analyzed the size and intensity for both antibody- and nanobody-revealed molecular arrangements from STED images. For an accurate comparison between the two types of probes, the signals were expressed as fold over the signal of single antibody complexes and nanobodies measured in similar STED images. The analyzed regions were separated in synaptic regions, strongly immunostained for Synaptophysin and non-synaptic regions, devoid of Synaptophysin signals. The analysis revealed that SNAP-25 nanobodies detected ~4-fold more epitopes within synapses and ~6-fold more epitopes outside of synapses (extra-synaptic) than SNAP-25 antibodies (Figure 7A). This suggests that SNAP-25 clusters/spots contain at least 10-80 molecules within synapses, and at least ~10 molecules outside of synapses (since the nanobodies do not necessarily detect all of the molecules, and therefore the number of nanobodies detected per cluster is only a minimal estimate for the number of molecules). For Syntaxin 1A, both antibody and nanobody immunostainings revealed ~20 molecules per cluster within synapses, while the nanobody revealed ~8 molecules per cluster outside of synapses, with only 1-2 revealed by the antibody (~4-fold difference; Figure 7B).

**Figure 7:**
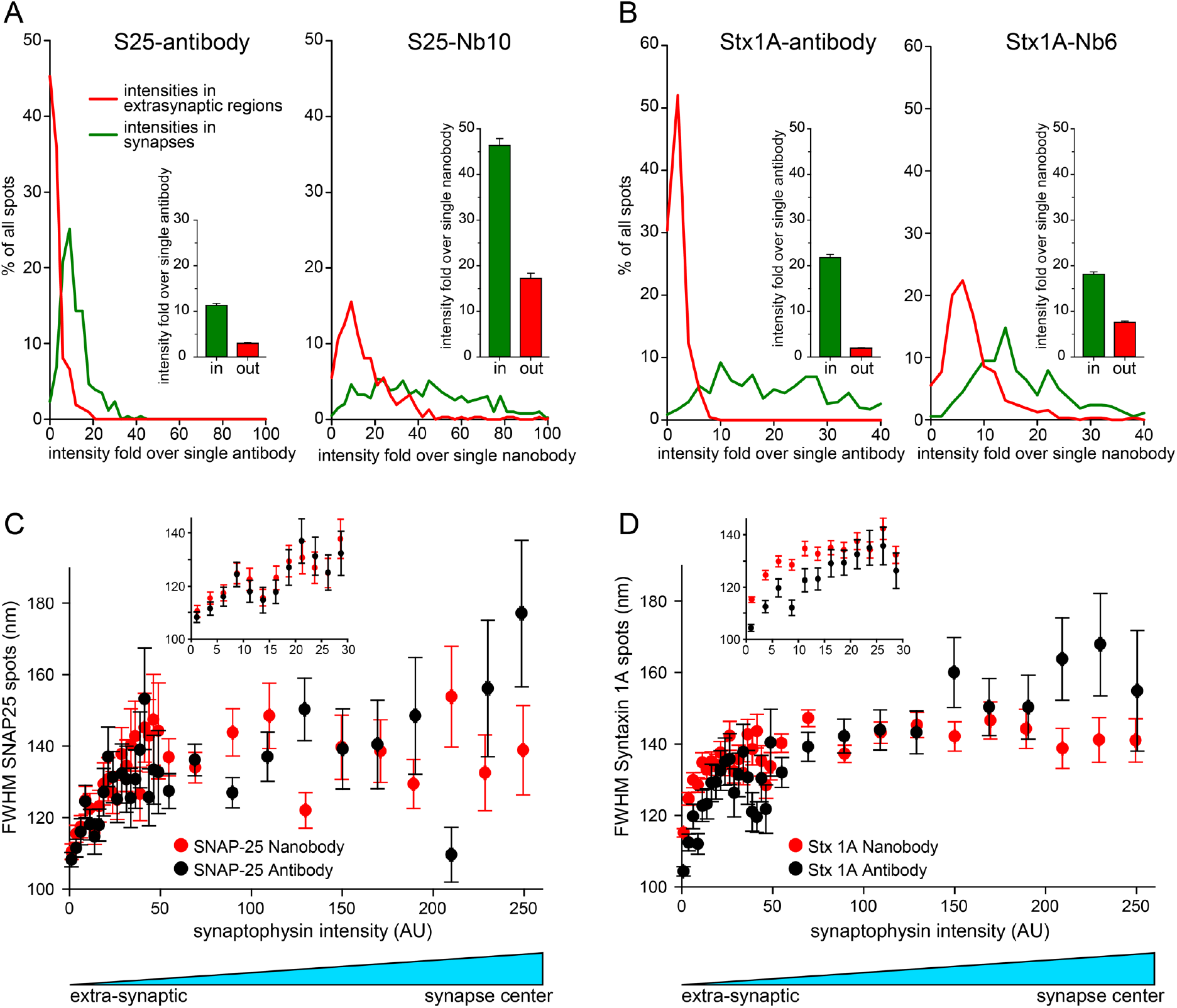
An analysis of the size and intensity of SNAP-25 and Syntaxin 1A clusters (spots) in hippocampal neurons. **(a)** The intensity of SNAP-25 spots is shown as intensity fold over single nanobody or antibody. The intensities were obtained by drawing line scans across 259 SNAP-25 clusters in synapses and 212 clusters outside of synapses for the antibody staining. For the nanobody staining, we analyzed 392 line scans in synapses and 309 line scans outside of synapses. Both within and outside of synapses, the antibody shows spots of a smaller intensity than the nanobody. The nanobody spots in synapses (green line in right panel) also show a large variability within the signal intensity. Bar graphs (insets) represent the mean of the intensity distributions with their associated SEM. **(b)** Same analysis as in **(a)** for Syntaxin 1A. We analyzed 348 line-scans in synapses and 227 outside synapses for antibodies. 378 line-scans were analyzed in synapses and 326 outside synapses for nanobodies. Interestingly, the nanobody spots in synapses (green line in right panel) show a far more limited variability than for SNAP-25 (note the difference in units for the X-axis of SNAP-25 and Syntaxin 1A analyses). **(c-d)** Analysis of spot size for SNAP-25 or Syntaxin 1A revealed by nanobody (red) or antibody (black), relative to the Synaptophysin intensity (very low signal is extra-synaptic, highest signal is the center of the synapse). The insets present the extra-synaptic areas at higher zoom in the regions of low Synaptophysin intensity. No obvious difference can be observed for SNAP-25 **(c)**, but the Syntaxin 1A spots (clusters) are larger for the nanobody in extra-synaptic areas (p = 0.000082, paired t-test). The analysis was performed on the following number of automatically identified spots: 10,839 spots from seven independent experiments for SNAP-25 nanobodies; 8,546 spots from seven independent experiments for SNAP-25 antibodies; 35,609 spots from 20 independent experiments for Syntaxin 1A nanobodies; 12,468 spots from seven independent experiments for Syntaxin 1A antibodies. The graphs show bi-directional scatter plots, plus SEM, after binning the spots according to Synaptophysin intensity. Note that for the Synaptophysin staining the error bars in the horizontal direction are smaller than the symbol sizes.

This analysis reinforces the concept that the two molecules form clusters but suggests that they are also prominent outside of synapses. At the same time, an analysis of the size (diameter) of the clusters revealed that they averaged ~140 nm in synapses (Figure 7C, 7D). Outside of synapses, the cluster size was smaller, down to ~110 nm. Syntaxin 1A nanobodies detected larger clusters than antibodies outside of synapses, presumably because the limited staining provided by antibodies outside synapses was not sufficient to resolve clusters reliably in this region.

In order to further investigate the extra-synaptic populations of SNAP-25 and Syntaxin 1A, we decided to follow their distribution upon electrical stimulation (60 seconds at 20 Hz). The localization of SNAP-25 and Syntaxin 1A at the vicinity of synapses (determined by the Synaptophysin staining as above) was investigated by STED microscopy. The samples were fixed and stained without stimulation (as a control), immediately after the electrical stimulation, or after five minutes of recovery at 37 °C. The results suggest that the population of SNAP-25 located outside of synapses does not show a detectable net movement upon stimulation. In contrast, the extra-synaptic population of Syntaxin 1A had relocated to synapses upon stimulation. This population then recovered and distributed again to the extra-synaptic regions within five minutes after stimulation (Figure 8). Overall, this does not only confirm the specificity of our probes, but also provides completely new evidence for the recruitment of a SNARE molecule (Syntaxin 1A) into synapses during strong stimulation.

**Figure 8:**
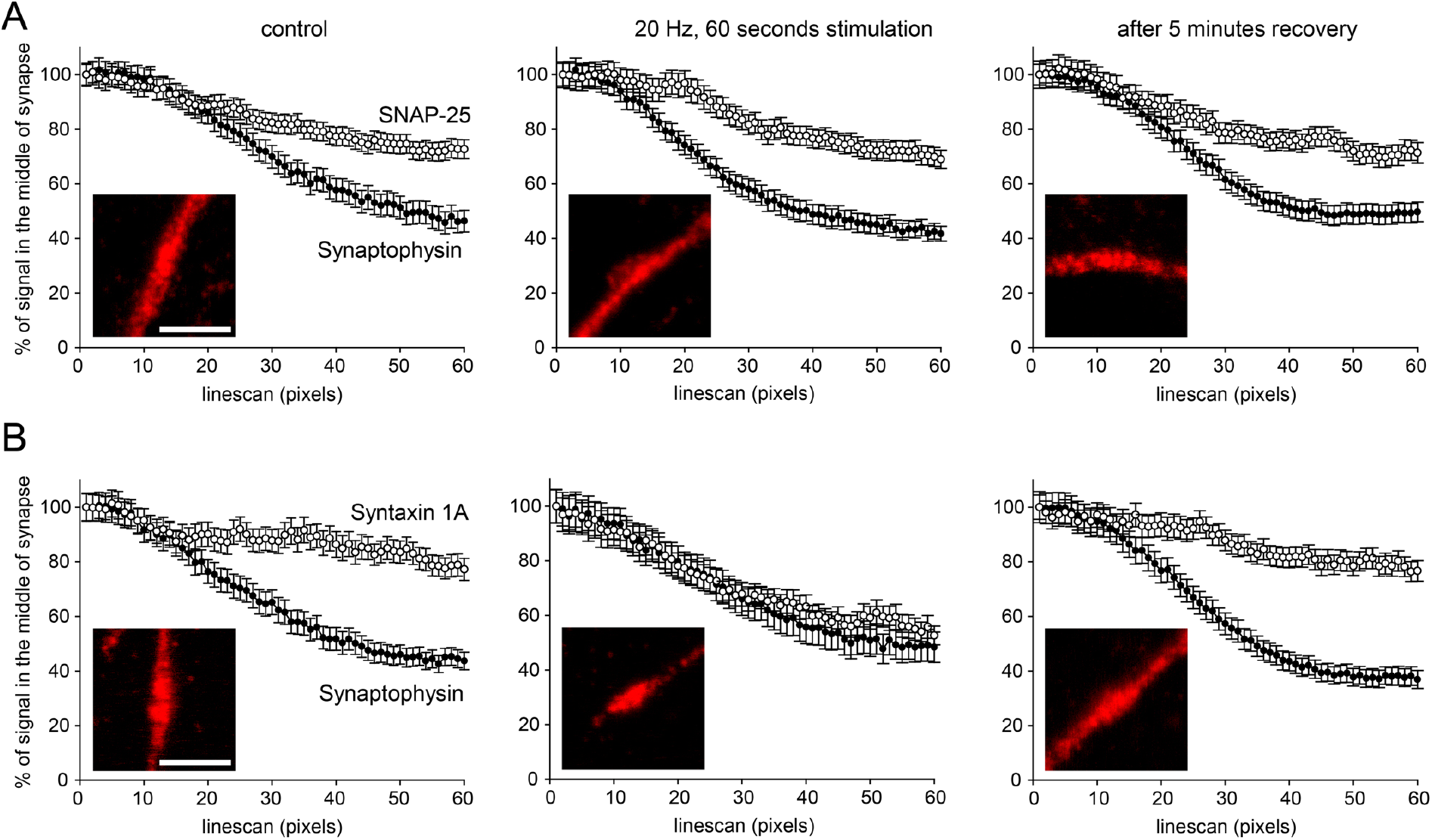
Investigation of the extra-synaptic populations of SNAP-25 and Syntaxin 1A in hippocampal neurons upon electrical stimulation. Cultured neurons were co-stained with nanobodies directly coupled to Atto647N and with anti-Synaptophysin antibodies (detected with a secondary coupled to Alexa488) to identify synaptic regions. **(A)** STED images of the nanobody immunostainings were analyzed. The distribution of extra-synaptic SNAP-25 remains unchanged upon stimulation at 20 Hertz for 60 seconds, resembling the pattern previously observed in Figure 5. For control, stimulation and recovery conditions, 356, 388 and 424 synapse line profiles were analyzed from three independent experiments. **(B)** Upon stimulation, the population of Syntaxin 1A that is outside of synapses relocates into neighboring synapses. This population recovers if the neurons are further incubated for five minutes at 37 °C after stimulation. For control, stimulation and recovery conditions, 288, 144 and 280 synapses were analyzed from three independent experiments. The exemplary images show the nanobody signals in regions centered on synapses (determined by Synaptophysin staining). The scale bars represent 1 μm.

## Discussion

Detecting molecules of interest by affinity reagents has been an invaluable tool in the last decades of biomedical research. A rapidly developing field for such tools is the development of single domain antibodies, or nanobodies, derived from camelids. Nanobodies have several advantages when compared to conventional immunoglobulins.^15, 33^ For example, nanobodies can be produced recombinantly in expression systems like *E. coli*, minimizing the use of animals and increasing the scientific reproducibility, as batch effects are eliminated.^16, 34^ Additionally, nanobodies have caught the attention of the fluorescence super-resolution microscopy field due to their small size (~3 nm, Figure 1), which results in minimal displacement errors between the location of the fluorophore and that of the target.^13,14,35^ We therefore made efforts to generate nanobodies binding two of the major SNARE proteins involved in neuronal exocytosis: SNAP-25 and Syntaxin 1A. Our objective was to obtain highly specific nanobodies able to detect the endogenous proteins in a more precise manner than possible with classical antibodies. The selected nanobodies displayed a high specificity and affinity to their intended targets and revealed a population of SNAP-25 and Syntaxin 1A in hippocampal neurons that could not be detected by conventional antibodies (Figure 5). Furthermore, the Syntaxin 1A nanobodies provided evidence for a high, stimulation-dependent mobility for the extra-synaptic population of Syntaxin 1A, which implies that these molecules presumably play a role in exocytosis upon prolonged stimulation.

### Selection and characterization of the nanobodies

During two rounds of phage-display screening, we selected for binders that should be able to support stringent and prolonged washing conditions, as required to perform background-free immunostainings. The monovalent nanobodies ultimately selected bind with dissociation constants (K_D_) in the low nM range (Supplementary Figure 2) and exhibit a very high specificity, even if challenged against other conserved homologs or isoforms (Supplementary Figure 3).

Both nanobodies clearly revealed only one band in Western Blots, regardless whether the SNAP-25 and Syntaxin 1A targets were produced recombinantly in bacteria or mammalian cells (HEK293), or endogenously in rat brain or primary neuron lysates (Figure 3C, 3D). This implies that they only detect single targets and that these are the intended neuronal exocytosis SNAREs, as no signals could be detected in non-neuronal tissues (Supplementary Figure 5).

### The nanobodies reveal target populations that are more poorly identified by the antibodies

A surprising finding was that the nanobodies showed a similar behavior to antibodies in immunostainings, but revealed a much larger population of intracellular (perinuclear) signals in PC12 cells (Figure 3A). Both SNAP-25 and Syntaxin 1A are present in large copy numbers in PC12 cells,^36^ which implies that they need to be produced often and hence that they should be evident in the ER and Golgi membranes of the cells. Moreover, SNAP-25 appears to be highly accumulated in the Golgi of PC12 cells, because most of the activity of DHHC palmitoyl transferases, which are responsible for the membrane association of SNAP-25, takes place at the cytosolic surface of the Golgi.^31^ This suggests that the signals revealed by the nanobodies in the perinuclear regions are not artifactual, especially in view of the fact that the same behavior could be analyzed in COS-7 cells expressing EGFP chimeras of the proteins (Figure 4B-4D).

At the same time, this analysis demonstrated that the nanobodies can detect populations of epitopes that are largely missed by the antibodies. We assumed initially that this effect would be most evident in regions of high antibody staining intensity. Such regions presumably contain high levels of epitopes and we argued that the antibodies may not be able to reveal all of them due to steric hindrance, while the smaller nanobodies would be more efficient. The effect of steric hindrance was also striking in preparations of brain slices (Supplementary Figure 8) and appeared to also take place in COS-7 cells overexpressing the proteins (Figure 4B). Hence this argument seems plausible for cases in which protein crowding and/or penetration depth limit the antibodies (size 10-15 nm) more than the nanobodies (size 2-3 nm).

However, this argument does not seem to apply easily to the extra-synaptic regions of the hippocampal cultures. The molecules from these regions are revealed especially poorly by antibodies, although based on numerous investigations of axonal membrane morphology and apparent protein and lipid labeling in electron microscopy, we cannot argue that such regions are necessarily more crowded than the synaptic ones.^37^ A number of other effects probably limit the antibody ability to reveal the targets in these areas.

First, nanobodies rely on only one binding pocket, while the antibodies are divalent. The strong binding of antibodies to their targets depends on an avidity effect. Their probability to stay bound to targets is higher than for monovalent probes, because when one binding pocket unbinds from the target, the other is probabilistically still bound, thus strongly increasing the probability that the antibodies remain bound. However, this effect has an important downside. It only takes place in areas where the target is abundant, and both pockets can engage in target binding simultaneously. In areas with lower target densities, as outside synapses, the avidity effect is eliminated, and the antibodies have a much higher chance to be washed away.

Second, the nanobodies bind their epitopes stoichiometrically (one nanobody per target molecule and one fluorophore per nanobody), but this relation is known to be particularly heterogeneous for primary and secondary antibody detection systems. Both of the antibodies are divalent and typically polyclonal secondary antibodies are used. Thus, a primary antibody may be bound by multiple secondary antibodies. The avidity effect described in the previous paragraph functions for polyclonal antibodies as well and thus they become stabilized in regions of high density of primary antibodies. Such regions therefore may present disproportionally strong signals.

Third, fixation by PFA, which is typical for most immunostainings, leaves a large fraction of the targets unfixed and therefore mobile.^38, 39^ Single target molecules, attached to a single antibody binding pocket, can therefore diffuse until they reach areas where other target molecules are present. Here the second antibody binding pocket can become bound to a second target molecule. The larger molecular arrangement of two target molecules and one antibody is less mobile and is more likely to remain in the area of high target density than to return to the initial low-density area. It is then stabilized further by the secondary antibodies, and again contributes to disproportionally higher signals within the areas where the target density was higher.

Finally, it may be that the antibody epitope is masked by an interacting partner or buried in a different fold conformation in this extra-synaptic population of SNAP-25 and Syntaxin 1A. All these effects were not measured directly in this work, and therefore their discussion is to some extent speculative.

### The nanobodies provided new information on the distribution of SNAP-25 and Syntaxin 1A on primary hippocampal neurons

The two nanobodies binding SNAP-25 and Syntaxin 1A provided a number of observations that change, at least to some extent, our view on the membrane organization of these two molecules. As indicated in the introduction, one of the most important features of the organization of both SNAP-25 and Syntaxin 1A has been their presence in clearly distinguishable clusters, which have been some of the first objects of study by super-resolution microscopy.^7, 8^ These clusters are detected with nanobodies as expected, since GFP-tagged versions of the molecules also cluster in membranes.^7, 40^ At the same time, some studies described membranes containing virtually only large and bright clusters, which are clearly separated in membranes by cluster-free areas.^9^ However, this may as well be an artefact of the antibody stainings,^17^ which yields a disproportional fluorescent signal as discussed above. In fact, the clusters revealed by the nanobodies seem to be especially heterogeneous for SNAP-25 (Figure 7A), with clusters varying by ~8-fold in brightness being found with similar probabilities. Importantly, this is not the case for Syntaxin 1A, for which a previous study emphasized the fact that inherent structural features of the molecule should limit the size of the clusters that can be formed.^8^ We indeed found that the size of these clusters was much more homogenous than the one of SNAP-25 clusters (Figure 7B).

Two additional important effects became evident using nanobodies for stainings. First, while the SNAP-25 and Syntaxin 1A clusters rarely overlapped to a significant effect in the literature, they tend to do so when immunostained by nanobodies (Figure 6). This confirms the observation that both SNAP-25 and Syntaxin 1A are located to the same areas of membrane protein clusters or islands.^41^ Second, the existence of a substantial extra-synaptic population of these molecules changes the current perspective on their functional organization: SNAP-25 and Syntaxin 1A are also axonal, rather than purely synaptic proteins. Their special features, ranging from cluster formation to peculiar membrane interactions^42^ or extremely rapid axonal transport^43^ should therefore be discussed in this perspective. For example, this implies that the SNAP-25 and Syntaxin 1A localization is unlikely to play a major role in defining exocytosis, with the locations of other elements such as including SNARE-regulating proteins^44^ being more important in this respect.^5^

The described differing patterns observed with nanobodies and antibodies are presumably not limited to the distribution of SNAP-25 and Syntaxin 1A and thus we suggest that nanobody development is also desirable for other target proteins. We conclude that small, monovalent affinity tools do not only have the potential of providing higher quality super-resolution images,^13, 14^ but may also reveal quantitative and qualitatively different features, especially by their linear signal-to-target stoichiometry and by revealing target molecules that are present in low copy numbers or buried in crowded regions.

## Methods

### Protein expression and purification

Full antigens and their truncated form (Figure 1 and 2) were produced by recombinant expression in *E. coli* NEB Express strain (New England Biolabs, Ipswich, MA, USA). The expression vector was derived from the LacO-pQLinkN-construct^45^ containing a N-terminal Histidine-tag and a bdSUMO-domain fused to the protein of interest to increase protein solubility and allow cleavage with bdSUMO protease.^46^ A Twin-Strep-Tag (IBA GmbH, Göttingen, Germany) was fused to the C-terminus of the protein for affinity purification. Protein expression was induced by adding IPTG to a final concentration of 1 mM. Cultures were grown in Terrific Broth (TB) medium (Sigma-Aldrich, St. Louis, MO, USA) overnight at 30 °C while shook at 120 rpm.

Similarly, nanobodies were expressed in *E. coli* SHuffle Express bacteria (New England Biolabs) at 25 °C overnight, 120 rpm. The next day, bacteria were harvested by centrifugation for 20 minutes at 3200 x g. The pellet was resuspended in lysis buffer containing 50 mM HEPES, 150 mM NaCl, 1 mM EDTA, 1 mM PMSF, 500 μg/ml lysozyme and 100 μg/ml DNaseI (all from Sigma-Aldrich) at pH 8.0. After incubation on ice for 20 minutes, bacteria were lysed by sonication with a Branson DigitalSonifier (Branson Ultrasonics, S. Louis, MO, USA) applying five times five pulses at 95% power. Subsequently, cell debris were removed by centrifugation at 11,000 x g for >1 hour at 4 °C. The crude lysate was filtered through a 0.45 μm syringe top filter (Sartorius, Göttingen, Germany) purified on an ÄKTApure25 HPLC system using StrepTrap HP or HisTrap HP columns (GE Healthcare, Little Chalfont, United Kingdom). After competitive elution from the column using binding buffer supplemented with either 500 mM ultrapure imidazole (AppliChem, Darmstadt, Germany) or 7.5 mM desthiobiotin (Sigma-Aldrich), the bdSUMO-domain was cleaved off by adding self-produced bdSUMO-protease. The cleaved fragment (His-Tag-bdSUMO) was removed using cOmplete His-Tag purification resin (Roche, Basel, Switzerland). The purity of the proteins was evaluated in a Coomassie stained PAGE (>95% pure) and protein concentration was finally measured by Novagen BCA-assay (Merck, Darmstadt, Germany). Proteins were snap-frozen in liquid nitrogen and stored at −80 °C.

### Molecular cloning

Cloning of constructs for protein expression was done according to Gibson *et al*^47^ Vector and insert for assembly were combined in a concentration of 15 fmol each in reaction buffer and incubated at 48°C for 30 minutes. 1 μl of the Gibson reaction was used to transform competent *E. coli* DH5α™ bacteria (Thermo Fisher Scientific, Waltham, MA, USA) which were subsequently grown over night on LB-agar plates supplemented with 50 μg/ml carbenicillin or kanamycin. Individual colonies were grown overnight in 5 ml LB medium including the corresponding antibiotics and plasmids were isolated using GeneJET™ plasmid purification kit (Thermo Fisher Scientific) for subsequent sequencing to confirming the plasmid Identity (SeqLab, Göttingen, Germany).

### Cell line culture

COS-7 cells were cultured as described before^48^ in high glucose (4.5 g/l) Dulbecco’s Modified Eagle’s Medium (DMEM) supplemented with 4% glutamine, 10% fetal calf serum (FCS) and 100 U/ml penicillin-streptomycin. PC12 cells were cultured in DMEM supplemented with 4 mM L-glutamine, 10% Horse Serum (HS), 5% FCS and 100 U/ml penicillin-streptomycin as described before.^49^ Cells were finally seeded on coverslips pre-cleaned with 1 M NaOH, followed by 1 M HCl and finally by 100% ethanol. After thoroughly washing with sterile water, coverslips were coated with 0.1 mg/ml poly-L-lysine (PLL, Sigma-Aldrich). 12 to 16 hours after incubation in a humidified incubator at 37 °C and 5% CO2, cells were transfected using Lipofectamine2000^®^ (Thermo Fisher Scientific). Typically, cells were used for immunostaining experiments the day after transfection (16-20 h).

### Primary neuron culture

Primary hippocampal neurons were cultured as described before.^50^ Alternatively, neurons were cultured according to Kaech and Banker^51^ to obtain low density neuron cultures for immunofluorescence experiments. Briefly, glia cells were prepared from cortex and seeded directly into 12-well cell culture plates in Minimum Essential Medium (MEM, Sigma-Aldrich) supplemented with 10% HS, 0.6% glucose, 1 mM L-glutamine and 100 U/ml penicillin-streptomycin. After four days, primary neurons were seeded on glass coverslips which have been pre-cleaned with nitric acid and coated with 1 mg/ml PLL before. These neuron containing coverslips were incubated onto the glia cells containing wells, coverslips were placed upside down, so the cells (glia and neurons) are facing each other. A few small paraffin dots on the coverslips allowed spatial separation of the two cultures. The culture medium was replaced by neuronal maintenance medium as described by Kaech *et al.^51^* Neurons with 12-18 days in vitro (DIV) were finally used for immunostainings.

### Nanobody library construction

An alpaca *(Vicugna pacos)* was repeatedly immunized with 300-500 μg of both recombinant purified SNAP-25 and Syntaxin 1A suspended in incomplete Freud’s adjuvant (performed by Preclinics GmbH, Potsdam). A total of six injections were given to the animal at an interval of one week. Two days after the final injection, 50 ml of blood were taken and peripheral white blood cell were isolated by Ficoll-gradient centrifugation. Total RNA from that preparation was extracted using RNeasy purification kit (Qiagen, Hilden, Germany). Subsequently, cDNA was synthetized from extracted total RNA using the SuperScript IV Reverse Transcriptase (Thermo Fisher Scientific) and primers specifically aligning to the conserved IgG framework. The protocol for the amplification of nanobody sequences by nested PCR was adapted from Olichon and de Marco.^52^ First, the overall nanobody repertoire was amplified using the universal primers GTCCTGGCTGCTCTTCTACAAGG (CALL1) and GGTACGTGCTGTTGAACTGTTCC (CALL2) for ten cycles. Next, the nanobody sequences were further amplified using a mixture of primers specific for IgG2 and IgG3 subtypes aligning to the framework and hinge region for eight cycles in total. Forward primer sequence: TCTGGTGATGCATCTGACAGCGAGGTGCAGCTGSWGGAGTCTGG Reverse primer sequences: GTTTTCCCCAGTGGATCCAGAACTAWTAGGGTCTTCGCTGTGGTGC and GTTTTCCCCAGTGGATCCAGAAGTTTGTGGTTTTGGTGTCTTGGG. The primer overhangs (highlighted in gray) were used for subsequent Gibson cloning into a phagemid vector as described above. For construction of a phage display library, a modified version of the pHen2 phagemid kindly provided by Dr. Frank Perez was used.^53^ A FLAG-tag was included in the phagemid backbone to allow detection of the expressed candidates by anti-FLAG antibodies. The ligated vectors were transformed into electrocompetent TG-1 *E. coli* (Lucigen, Middletown, WI, USA) using 50 individual electroporations to maximize library diversity. Subsequently, all transformation reactions were pooled and distributed on 20 square (25 x 25 cm) LB-agar plates containing 50 μg/ml carbenicillin and incubated over night at 37 °C. Dilution series of the transformations were plated to determine overall library size. Next day, bacteria were scraped off the plates using LB medium and supplemented with glycerol for storage at −80 °C. The overall library size was found to be ~4 x 10^7^ colony forming units (cfu).

### Phage display and nanobody selection

The phage display screening was adapted from a protocol by Olichon *et al*.^54^ and Lee *et al*.^55^ Briefly, an aliquot of the library was diluted into 500 ml 2xYT medium supplemented with 4% glucose and 50 μg/ml carbenicillin for growing at 37 °C, 120 rpm. When OD600 reached 0.5, MK-13 helper phages (#N0315S, New England Biolabs) or M13 K07ΔpIII hyperphages (PROGEN Biotechnik, Heidelberg, Germany) were used to infect the culture. Phages were produced over night at 30 °C at 120 rpm in 500 ml 2xYT medium, supplemented with 50 μg/ml of both carbenicillin and kanamycin. Next day, the culture was pelleted and phages were precipitated from the supernatant by adding polyethyleneglycol (PEG) and NaCl to final concentrations of 5% and 500 mM, respectively. The final phage titer was determined by measuring the OD260 using the empirical formula given by Lee *et al.^55^* Antigens were immobilized to MagStrep “type3” XT beads (IBA GmbH) via a C-terminal Twin-Strep-Tag fused to the protein. Phages were incubated with the immobilized antigen for 1 hour at room temperature. Afterwards, a total of 10 washes with PBS were performed for at least ten minutes per washing step. Retained phages were eluted using Strep-Tactin Biotin Elution Buffer (IBA GmbH) for 30 minutes at room temperature. The eluted phages were subsequently used to infect 50 ml of *E. coli* TG-1 bacteria (Lucigen) culture grown to OD600 ^Ä^ 0.5. After incubation for 1 hour at 37 °C, the culture was pelleted and resuspended in 1-2 ml of 2xYT medium. The resuspension was plated on 2xYT-agar plates supplemented with 50 μg/ml carbenicillin to select infected bacteria. The next day, colonies were scraped off the agar plates, diluted into 2xYT medium and grown as described above for a new round of biopanning. Typically, 2 panning rounds were performed for each screening procedure, successively increasing the stringency of binding and washing conditions in each round. For the initial panning, M13 K07ΔpIII hyperphages (PROGEN Biotechnik) were used instead of MK-13 helper phages to increase the amount of initially retained nanobodies/phages. After the final biopanning round, individual colonies were picked and grown in 96-well plates for a monoclonal phage-ELISA. MK-13 helper phages were added to the wells to produce single-clone phages overnight. The antigen was immobilized on Nunc MaxiSorp flat-bottom 96-well plates (Thermo Fisher Scientific) and blocked 1 hour with 5% milk powder in PBS-T. The phage supernatant of infected colonies was mixed the immobilized antigen and incubated for 1 h. After 3 washes with PBS for ten minutes, bound phages were detected with an HRP-coupled antibody directed against the phage pVIII-protein (clone B62-FE2, Abcam, Cambridge, United Kingdom). Positive binding was detected using 1-Step Ultra TMB-ELISA substrate solution converted by the HRP (Thermo Fisher Scientific). The reaction was stopped by adding 2 M H2SO4 followed by readout of the absorbance at 430 nm in a Cytation-3 Multi-Mode Reader (BioTek Instruments, Winooski, VT, USAUSA). Binding of individual candidates was considered positive if the ratio of the read signal was at least ten fold over background and negative controls. Finally, cultures from the positively tested wells were grown for sequencing of phagemids (SeqLab) and further validation.

### Immunoblotting (dot-blots)

For simple validation of target binding (Supplementary Figure 1B), 1 μg of purified antigen was spotted on a nitrocellulose membrane (Amersham, Sigma-Aldrich). After blocking for 1 hour with 5% milk in PBS-T, the membrane was incubated with phage-containing supernatant from monoclonal phage ELISA in 2.5% milk/PBS-T for 1 hour at room temperature. Bound nanobodies were detected using an anti-DDDDK-tag antibody coupled to DyLight^®^650 (clone M2, Abcam). After washing with PBS-T, membranes were imaged in an Amersham Imager 600 (GE Healthcare). For specificity and epitope mapping analysis in Figure 2, dot blots were performed as mentioned above, but detection was performed using directly conjugated nanobodies to Atto647N fluorophore and read in the Amersham Imager 600 (GE Healthcare). Homolog proteins of SNAP-25 and Syntaxin 1A were all purchased from OriGene, Rockville, MD, USA and truncated versions were clone and produced as explained above.

### Tissue isolation

To confirm the binding specificity of the nanobody candidates, different animal tissues were isolated from adult mice. Immediately after dissection, the tissues were homogenized in ice-cold PBS supplemented with 1 mM EDTA and protease inhibitor cocktail (Sigma-Aldrich). The tissue was homogenized with a motor-driven glass-Teflon homogenizer (Omnilab, Bremen, Germany) at 900 rpm, 30 strokes and snap-frozen in liquid nitrogen. Total protein concentration was determined by Novagen BCA-assay (Merck).

### Cell and brain lysate preparation

HEK293-FT cells transiently expressing SNAP-25 or Syntaxin 1A or primary cultured hippocampal neurons were suspended in ice-cold lysis buffer composed of 50 mM Tris/HCl, pH 7.5, 150 mM NaCl, 2 mM EDTA, 0.5% (v/v) IGEPAL and 0.5% (w/v) Sodium deoxyclolate supplemented with 1 μg/ml DNAse I, 10 μg/ml aprotinin, 10 μm/ml leupeptin, 1 μg/ml pepstatin A, 100 μM PMSF (all from Sigma-Aldrich) and 0.1x Protease Inhibitor Cocktail (Roche). Suspended cells were incubated on ice for 45 minutes followed by five sonication pulses of three seconds (Branson Ultrasonics). After incubation on ice for another 15 minutes, cell debris was removed by centrifugation for 45 minutes at 4 °C, 16,000 x g and the supernatant containing the soluble cell lysate was snap-frozen in liquid nitrogen. Whole brain lysate was prepared accordingly after grinding the tissue on ice with a pellet mixer (VWR, Radnor, PA, USA) in ice-cold lysis buffer.

### Gel electrophoresis and Western blotting

Proteins were analyzed by SDS-PAGE according to Schagger and von Jagow^56^ on a 10% denaturating Tris/Tricine polyacrylamide gel. The Mini-Protean Tetra Cell System (BioRad, Hercules, CA, USA) was used to run the gel a discontinuous buffer system at 90 volts for 120 minutes. Proteins bands were visualized by staining overnight in Coomassie Brilliant Blue-250 staining solution. Alternatively, polyacrylamide gels were blotted onto a nitrocellulose membrane in 50 mM Tris/HCl, 192 mM glycine, 20% methanol and 0.04% SDS. The membrane was blocked for 1 hour with 5% milk powder in PBS-T, and subsequently incubated with the fluorescently labeled nanobody at a concentration of 25 nM in 5% milk/PBS-T. After washing with PBS, membranes were imaged in an Amersham Imager 600 (GE Healthcare). For Supp. Fig. 5C and 5D, loading controls were performed by incubating the membrane with mouse anti pan-Actin antibody (NB600-535; Novus Biologicals) pre-incubated with an excess of FluoTag-X2 anti Mouse IgG IRdye CW800 (N1202; NanoTag Biotechnologies GmbH). Images were acquired using a LI-COR Odyssey Clx scanner.

### Fluorescent labeling of nanobody candidates

Purified S25-Nb10 and Stx1A-Nb6 both with an ectopic C-terminal cysteine was incubated on ice with 20-fold molar excess of TCEP for 30 minutes to open intermolecular disulfide bonds. The reduced nanobody was desalted into PBS pH 7.4 with a NAP-5 column (GE Healthcare) to remove unreacted TCEP. Subsequently, a maleimide-functionalized dye dissolved in anhydrous DMSO was added to the nanobody in 4-6 fold molar excess. The coupling reaction was stirred for two hours on ice shielded from light. Free dye was removed with a Superdex 75 gel filtration column (GE Healthcare). The degree of labeling (DOL) was calculated from the extinction coefficients and the absorbance of the dye and the protein. Only conjugates with a DOL >90% were used for immunostainings. After determining their concentrations and DOL, labeled nanobodies were brought to a 50% glycerol solution and stored at −20 °C. The nanobodies S25-Nb10 and the Stx1A-Nb6 have been licensed from the University of Göttingen Medical Center to NanoTag Biotechnologies GmbH to make them commercially available.

### Affinity measurements

To determine the dissociation constant (KD) of selected nanobodies we used microscale thermophoresis with a Monolith NT.115Pico Instrument (NanoTemper, Munich, Germany).

Fluorescently labeled nanobody was diluted into PBS containing 0.05% Tween-20, mixed with a dilution series of the ligand antigen and transferred into Premium Coated Capillaries (NanoTemper) as suggested by the manufacturer. KD-values were extracted from at least three independent experiments, using the Affinity Analysis software from NanoTemper.

### Immunostaining

Cells grown on PLL coated coverslips were fixed with 4% paraformaldehyde (PFA) prepared in PBS pH 7.4 for 45 minutes at room temperature. Remaining unreacted PFA molecules were quenched with 100 mM glycin and 100 mM NH4Cl in PBS for 20 minutes. To facilitate epitope accessibility, cells were subsequently permeabilized with 0.1% Triton-X100 in PBS for ten minutes under slow orbital shaking. To avoid unspecific binding of probes, cells were blocked for >1h with 3% bovine serum albumin (BSA) in PBS filtered through 0.22 μm syringe-top filter (Sartorius). Primary mouse monoclonal against SNAP-25 and Syntaxin 1A (Synaptic Systems, clone 71.1 #111011 and clone 78.2 #110011 respectively) were chosen due to their presence in a high number of studies.^5,39,40,57–59^ Guinea Pig polyclonal against Synaptophysin 1 (Synaptic Systems, #101004) or Rabbit polyclonal against SNAP-25 (Synaptic Systems, #111002) and Syntaxin 1A (Synaptic Systems, #110302) were incubated with the cells as recommended (typically at 67 nM or 1:100 dilution from the stocks). Incubation was performed in PBS supplemented with 1.5% BSA for 1 hour at room temperature. Alternatively, in Figure 4A the monoclonal HPC-1 antibody (Abcam, #ab3265) directed against Syntaxin 1A was used at 1:100 dilution for stainings of PC12 cells. After three washing steps with PBS for 10 minutes each, samples were incubated with the following fluorescently labeled secondary antibodies: Goat anti guinea pig coupled to AlexaFluor488, Jackson ImmunoResearch, #706-545-148; donkey anti rabbit coupled to Cy3, Dianova, #715-165-152; goat anti rabbit coupled to Abberior-Star580, Abberior, #2-0012-005-8; goat anti mouse coupled to Atto647N, Rockland, #610-156-121 as described before.^50, 60^ Alternatively, nanobodies conjugated to Atto647N or Aberrior-Star580 were used at final concentrations of 25-50 nM. Control staining of neurons (Supplementary Figure 9B) was performed using 25 nM of FluoTag-X2 anti EGFP Atto647N (NanoTag Biotechnologies GmbH). Nuclear staining for COS-7 and PC12 cells were performed using

Hoechst staining solution (Thermo Fisher). After three washings in PBS, coverslips were mounted in Mowiöl (6 g glycerol, 6 ml deionized water, 12 ml 0.2 M Tris buffer pH 8.5, 2.4 g Mowiöl 4–88, Merck Millipore) and dried over night at 4 °C.

### Imaging validation of first candidates

Nanobodies were grouped into families based on their complementary domain region 3 (CDR3) described by *Maas et al*.^61^ One representative member of each family was sub-cloned into SHuffle Express *E. coli* (New England Biolabs) for nanobody expression as described above. The crude lysate of each clone was incubated on 4% PFA fixed COS-7 cells transfected with the antigen of interest fused to EGFP. After 1 h incubation and three times washing with PBS, the bound nanobody was detected using an anti-DDDDK-tag antibody coupled to DyLight650 (clone M2, Abcam). A colocalization of the fluorescent antibody signal with EGFP without significant background binding was considered to indicate a specifically bound nanobody (Supplementary Figure 1B). Sequences of those candidates were sub-cloned into an expression vector for direct coupling to a fluorophore.

### Brain slice preparation and staining

Brain slices were prepared on ice from adult (6-8 weeks old) Wistar rats, by perfusion with PBS to remove blood, followed by incubation with 4% PFA for 60 minutes. The brains were removed from the skull and incubated in 4% PFA at 4 °C overnight. On the following day, brains were transferred to a solution of PBS supplemented with 30% sucrose at 4 °C until they sank to the bottom of the solution, before snap-freezing and storing them at −80 °C until sectioning into 30-35 μm thick slices on a Leica CM1850 cryotome. For staining, brain slices were incubated with primary antibody or directly conjugated nanobody PBS containing 3% BSA overnight at 4 °C. After three washing steps of ten minutes each, antibody samples were incubated with fluorescently labeled secondary antibody for three hours. After washing once again three times as before, slices were mounted on glass slides in thiodiethanol (TDE) by gradually increasing the concentration of TDE up to 100%. Finally, the mounted slices were sealed using nail polish to avoid drying of the sample.

### Pulldown and Co-Immunoprecipitation (Co-IP)

Nanobodies were conjugated to maleimide functionalized SulfoLink Coupling Resin (Thermo Fisher Scientific) according to instructions of the manufacturer. Successful conjugation was confirmed by adding 10 nmol recombinant antigen to 200 μl of the nanobody-functionalized beads (50% slurry). After extensive washing with PBS, protein elution was performed by boiling the beads in 2x SDS loading dye for ten minutes. Evaluation of the pulldown was done by a Coomassie SDS-PAGE analysis of the input, flow-through the beads, washing and elution fractions. For pulldown of SNAP-25 and Syntaxin 1A from tissue, total brain lysate was prepared as described as above. 200 μl of 50% slurry Nanobody-functionalized beads were incubated with 50 μl whole brain homogenate for 1 hour on ice and washed three times with PBS in a self-casted spin MoBiCol column (MoBiTec, Göttingen, Germany). Elutions were carried out by boiling the beads in 2x SDS loading dye for 10 minutes followed by SDS-PAGE analysis and Western blotting of all fractions obtained in the process. Similarly, purified SNARE-complexes containing full-length SNAP-25, Syntaxin 1A and Synaptobrevin 2 were prepared as before^3^ and subjected to immunoprecipitation using nanobody-functionalized beads. The SNARE-proteins of the complex were blotted on a nitrocellulose membrane and detected by monoclonal antibodies as described above.

### Stimulation of primary hippocampal neurons

To investigate the function of the extra-synaptic populations of SNAP-25 and Syntaxin 1A, 14-16 days in vitro primary rat hippocampal neurons were prepared as described before.^50^ Cells were transferred into 37 °C prewarmed Tyrode buffer containing 124 mM NaCl, 5mM KCl, 2 mM CaCl2, 1 mM MgCl2, 30 mM glucose and 25 mM HEPES (pH 7.4) and stimulated for 60 seconds at 20 Hz using a platinum plate stimulator (8 mm between the plates; driven from a stimulator and a stimulus isolator from World Precision Instruments, Friedberg, Germany). Subsequently, neurons were directly fixed using 4% PFA or kept in warm Tyrode buffer for another 5 minutes to recover before fixation. Immunostainings were performed as described before using the S25-Nb10 and Stx1A-Nb6 conjugated to Atto647N in combination with the synaptophysin antibody to identify the synapses.

### Fluorescence imaging

Qualitative binding validation (Supplementary Figure 1B) was done by epifluorescence imaging using an Olympus IX71 microscope equipped with 0.75 NA/60x oil objective and an Olympus F-View II CCD camera (Olympus, Hamburg, Germany). Confocal image acquisition was performed using a TCS SP5 STED confocal microscope (Leica Microsystems, Wetzlar, Germany) with a 100x 1.4 N.A. HCX PL APO CS oil objective (Leica Microsystems). Multichannel confocal images were obtained using an argon laser at 488 nm and helium/neon lasers at 594 nm and 633 nm for AlexaFluor488, Cy3, and Atto647N, respectively. Fluorescent signal was detected using photomultipliers. For STED microscopy, an inverse 4-channel Expert Line easy3D STED setup (Abberior Instruments GmbH, Göttingen, Germany) was used. The setup was based on an Olympus IX83 microscope body equipped with a plan apochromat 100x 1.4 NA oil-immersion objective (Olympus). Fluorescence excitation lasers (Abberior Instruments GmbH) pulsed at 40 MHz were utilized for the excitation lines 561 nm (for Abberior Star580) and 640 nm (for Atto647N). For depletion of the fluorescence signal of the Star580 and Atto647N dyes, a 775 nm STED laser (Abberior Instruments GmbH) pulsed at 40 MHz with an output power of ~1.250 W was used. Fluorescence signal was detected using APD detectors (Abberior Instruments GmbH) in predefined channels. The operation of the setup and the recording of images were performed with the Imspector software, version 0.14 (Abberior Instruments GmbH).

### Software and image analysis

Gene sequences were analyzed using ApE v2.0.47 by M. Wayne Davis or CLC sequence viewer v7.6.1 (Qiagen). Phylogenetic trees were created using CLC sequence viewer and molecular models were created with PyMOL Molecular Graphics System v1.7.4.5.

Image analyses of immunofluorescence experiments and plotting of line profiles (Figure 4) were performed using Fiji.^62^ The analyses of neuronal immunostainings were performed using self-written Matlab routines (Mathworks Inc, Natick, MA, USA). The line scans from Figure 5 were analyzed as follows: lines were drawn manually over axonal stretches containing multiple synapses. The centers of the Synaptophysin (synapse) signals were determined automatically, using a center-of-mass routine and the lines were broken into individual scans originating in the synapse centers, running in both directions for up to 500 pixels. The individual scans were then collected, overlaid and averaged, for the results shown in Figure 5D, 5E and in Figure 8. For Figure 6, we analyzed the Pearson’s correlation coefficients between the fluorescence signals in the different channels, exclusively within the synaptic areas which were defined by the Synaptophysin immunostainings. The areas were determined by applying empirically determined thresholds in the Synaptophysin immunostainings. For Figure 7A, 7B, an experienced user drew line scans manually over several hundred evident protein clusters. The line scans were fitted with Lorentzian curves, and the cluster intensity (summed over the entire fit) and size (full width at half maximum, FWHM of the fit) were measured. These intensities were later expressed as fold over the intensity of single antibodies or nanobodies. For Figure 7C, 7D we performed a similar analysis, but in an automatic fashion, relying on an automated detection of the spots. This was performed by applying a band pass on the images, and eliminating all signals found under an empirically-derived threshold. The remaining spots were identified, and their intensity in all channels was measured; their sizes were determined by automatic fits, as above. This analysis, while less precise in the identification of spots than manually-drawn scans, has the advantage that it provides large numbers of spots, which enable the description of smaller effects on, for example, the spot size.

## Acknowledgments

We would like to thank Dr. Steffen Frey for providing reagents and support with Gibson assembly cloning, Dagmar Crzan for her help in preparation of rat brain slices and Dr. Katharina Seitz for helping us with the stimulation experiments. We also would like to thank Eugenio Fornasiero for his valuable input and proofreading of the text. This work was supported by the Deutsche Forschungsgemeinschaft (DFG) through Cluster of Excellence Nanoscale Microscopy and Molecular Physiology of the Brain (CNMPB) to FO. The work was further supported by grants to SOR from the European Research Council (ERC-2013-CoG NeuroMolAnatomy, 614765) and from the DFG (SFB1190/P09 and SFB1286/A03).

## Author contributions

FO conceived the project. MM, SOR and FO designed the experiments. AO provided valuable support for the phage display procedure. MM performed all experiments and acquired all microscopy data. MM, SOR and FO analyzed the data and wrote the manuscript.

